# Dissemination routes of the carbapenem resistance plasmid pOXA-48 in a hospital setting

**DOI:** 10.1101/2020.04.20.050476

**Authors:** Ricardo León-Sampedro, Javier DelaFuente, Cristina Díaz-Agero, Thomas Crellen, Patrick Musicha, Jerónimo Rodriguez-Beltran, Carmen de la Vega, Marta Hernández-García, R-GNOSIS WP5 Study Group, Nieves Lopez-Fresneña, Patricia Ruiz-Garbajosa, Rafael Canton, Ben S Cooper, Alvaro San Millan

## Abstract

Infections caused by carbapenemase-producing enterobacteria (CPE) are a major concern in clinical settings worldwide. Two fundamentally different processes shape the epidemiology of CPE in hospitals: the dissemination of CPE clones from patient to patient (between-patient transfer), and the transfer of carbapenemase-encoding plasmids between enterobacteria in the gut microbiota of individual patients (within-patient transfer). The relative contribution of each process to the overall dissemination of carbapenem resistance in hospitals remains poorly understood. Here, we used mechanistic models combining epidemiological data from more than 9,000 patients with whole genome sequence information from 250 enterobacteria clones to characterise the dissemination routes of the carbapenemase-encoding plasmid pOXA-48 in a hospital setting over a two-year period. Our results revealed frequent between-patient transmission of high-risk pOXA-48-carrying clones, mostly of *Klebsiella pneumoniae* and sporadically *Escherichia coli.* The results also identified pOXA-48 dissemination hotspots within the hospital, such as specific wards and individual rooms within wards. Using high-resolution plasmid sequence analysis, we uncovered the pervasive within-patient transfer of pOXA-48, suggesting that horizontal plasmid transfer occurs in the gut of virtually every colonised patient. The complex and multifaceted epidemiological scenario exposed by this study provides new insights for the development of intervention strategies to control the in-hospital spread of CPE.

## Introduction

Antibiotic resistance is one of the most concerning health challenges facing modern societies^1^. Antibiotic resistance is of particular concern in clinical settings, where resistant pathogens significantly increase the mortality rates of critically ill patients and the costs associated with infection management and control^1,2^. The spread of antibiotic resistance genes between bacteria commonly associated with nosocomial infections is mainly driven by the horizontal transfer of conjugative plasmids^3,4^. However, the frequency with which this occurs in the clinical settings and its importance for the dissemination of resistance at a local level remain poorly defined.

One of the most clinically relevant groups of nosocomial pathogens are enterobacteria that produce carbapenemases (β-lactamase enzymes able to degrade carbapenem antibiotics). Among carbapenemase-producing enterobacteria (CPE), clones of *Klebsiella pneumoniae* and *Escherichia coli* carrying plasmid-encoded carbapenemases pose the highest clinical threat^5^. Despite their clinical relevance, major gaps remain in our understanding of the epidemiology of CPE and of carbapenemase-encoding plasmids. Previous work has highlighted the importance of in-hospital CPE transmission from patient to patient^6,7^ (between-patient transfer). However, the dissemination and evolution of CPE in hospitals present an additional layer of complexity: the transfer of carbapenemase-encoding plasmids between enterobacteria clones in the gut microbiota of individual patients (within-patient transfer)^8,9^. Understanding the relative importance of between-patient and within-patient transfer is of central importance for understanding the epidemiology of CPE and informing intervention strategies to control the spread of carbapenem resistance in clinical settings.

One of the most frequent carbapenemases in enterobacteria is OXA-48^10^. OXA-48 was first described in a *K. pneumoniae* strain isolated in Turkey in 2001^11^ and is now distributed worldwide, with particularly high prevalence in North Africa, Middle Eastern countries, and Europe^10^. The *bla*_oχA-48_ gene is usually encoded in an IncL, broad-host-range conjugative plasmid called pOXA-48^8^ (Figure S1). This plasmid is frequently associated with *K. pneumoniae* high-risk clones^12^, such as the sequence types 11 (ST11), ST15, ST101, and ST405^6,13–15^, which are able to readily spread between hospitalized patients producing outbreaks of infections^16,17^. Previous epidemiological studies strongly suggested the possibility of within-patient pOXA-48 transfer^17–21^, indicating that pOXA-48 would be an ideal study system to investigate the nosocomial dissemination of carbapenem resistance.

In the present study, we examined the between-patient and within-patient transfer dynamics of plasmid pOXA-48 in a tertiary hospital over a two-year period. For our analysis, we used a large and well-characterized collection of pOXA-48-carrying enterobacteria generated at the *Hospital Universitario Ramon y Cajal* in Madrid as part of the European project R-GNOSIS (Resistance of Gram-Negative Organisms: Studying Intervention Strategies)^22,23^. Using statistical models and combining epidemiological data from more than 9,000 patients with whole-genome sequence information from 250 enterobacteria clones, we aimed to define pOXA-48 transfer dynamics at an unprecedented resolution. Specifically, we aimed to determine the relative contribution of between-patient and within-patient plasmid transfer in the epidemiology of pOXA-48, and to use these data to inform improved intervention strategies to control the spread of carbapenem resistance in hospitals.

## Results

### Patients colonised by pOXA-48-carrying enterobacteria in the hospital

During the R-GNOSIS project, hospitalised patients were periodically sampled to detect the presence of enterobacteria producing extended spectrum β-lactamases (ESBL) and carbapenemases in their gut microbiota (see methods). The study enrolled all patients admitted to two medical wards (gastroenterology and pneumology) and two surgical wards (neurosurgery and urology) in the hospital. The full details of the R-GNOSIS study in our hospital, including the study population and CPE characterization, have been previously reported by Hernandez-Garcia *et al*^18,23^. Briefly, from March 2014 to March 2016, 28,089 rectal swabs were collected from 9,275 patients, and 171 pOXA-48-carrying enterobacteria strains were isolated and characterised from 105 patients (Figure 1, Table S1). The proportion of patients who were found to be colonized with pOXA-48-carrying enterobacteria on at least one occasion during their hospital admission was 0.5% in urology (18/3,483), 1.3% in gastroenterology (33/2,591), and 1.5% both in neurosurgery (16/1,068) and pneumology (38/2,559), with the medical wards accounting for 68% of colonised patients (71/105, Figure 1A-C).

**Figure 1.**
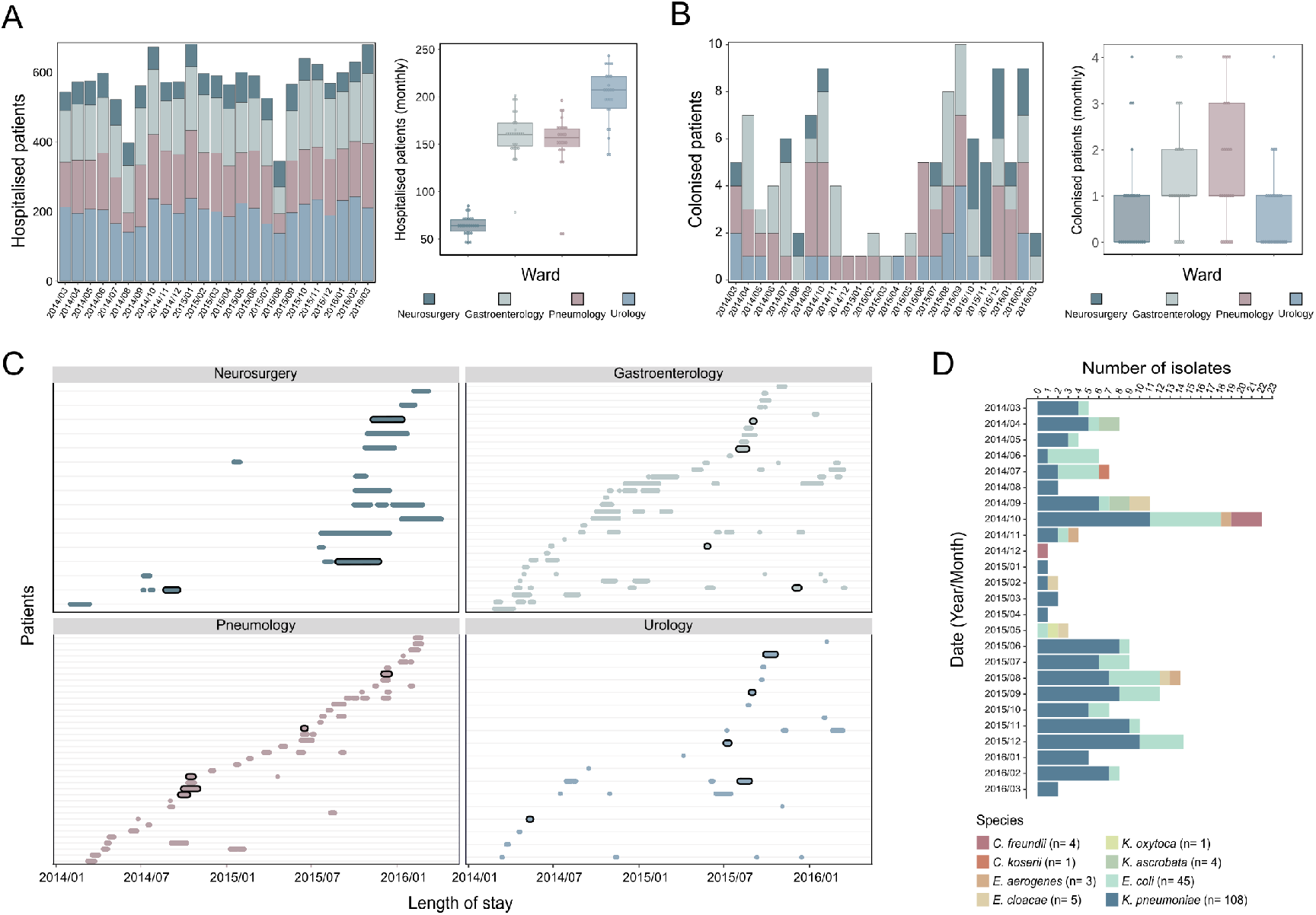
Study population, colonised patients, and pOXA-48-carrying enterobacteria. (A) Patients sampled during the R-GNOSIS study. The left panel shows the number of hospitalised patients in each ward over the 25-month study period. The right panel shows the distribution of hospitalised patients per ward by month as a boxplot. Horizontal lines inside boxes indicate median values, the upper and lower hinges correspond to the 25th and 75th percentiles, and whiskers extend to observations within 1.5 times the interquartile range. (B) Patients colonised by a pOXA-48-carrying enterobacteria during the study. The left panel represents the number of colonised patients in each ward over the 25-month study period. The right panel shows the distribution of colonised patients per ward by month as a boxplot. (C) Distribution of patients colonised by pOXA-48-carrying enterobacteria in the four wards under study over the 25-month study period. Each row represents a patient, and the colour segments represent the length of hospital stay (from admission to discharge). Black outlining of colour segments indicates patient co-colonisation with more than one pOXA-48-carrying species. (D) Enterobacteria isolates carrying pOXA-48 recovered from the patients during the 25 months of the study. The species of the pOXA-48-carrying isolates are colour-coded as indicated in the legend.

In line with previous reports^10^, *K. pneumoniae* was the most frequent pOXA-48-carrying species (n= 108). However, pOXA-48 was detected in an additional 7 enterobacterial species, with *E. coli* being the second most frequent carrier (n= 45, Figure 1D, Table S1). In several pOXA-48 carrying patients (18/105), there was cocolonisation of the gut microbiota with more than one species carrying the plasmid, suggestive of within-patient plasmid transfer events (Figure 1C).

### Using epidemiological data to analyse pOXA-48 transfer dynamics

To investigate how pOXA-48 spread in the hospital, we analysed the epidemiological data using a previously described model which enabled us to estimate the daily probability of a patient acquiring pOXA-48-carrying enterobacteria and quantify the effect of covariates on this probability (see methods and Supplementary Table 2)^24^. We performed this analysis independently for the two species with a large number of isolates, *K. pneumoniae* and *E. coli,* and we included two covariates in the model. The first covariate was the number of other patients colonized by pOXA48-carrying enterobacteria in the ward on the same day, which we expect to be positively associated with the daily risk of acquisition if between-patient bacterial transfer is important. The second covariate was known pre-existing intestinal colonisation of the patient by a pOXA48-carrying enterobacteria of a different species (*K. pneumoniae* or *E. coli).* If within-patient plasmid transfer is important (from *K. pneumoniae* to *E. coli* and *vice versa,* Figure 2), then we would expect this to also be positively associated with the daily risk of a patient acquiring pOXA-48-carrying enterobacteria. We considered different transmission models including and excluding these covariates and performed model comparisons using the widely applicable information criterion (WAIC, Supplementary Table 2). The model that best fitted our data was the one including both covariates and permitting the transmission parameter β to vary by ward (see Supplementary Table 3 for daily probability values and methods for details).

**Figure 2.**
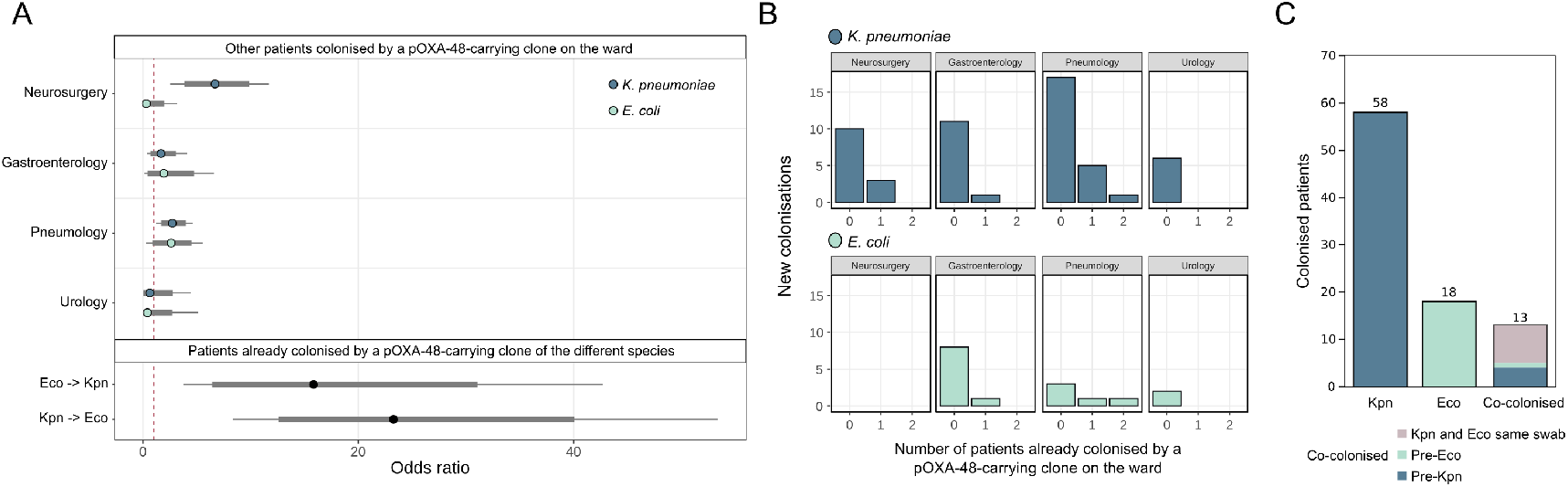
Acquisition of pOXA-48-carrying enterobacteria by hospitalised patients. (A) Posterior distribution of odds ratio for the daily risk of colonisation with a pOXA-48-carrying *K. pneumoniae* or *E. coli.* Two covariates were included. The first is the presence of other patients colonised by a pOXA-48-carrying clone on the ward, (upper part, stratified by ward). If between-patient transfer of the plasmid is important, we expect to see a positive association (odds ratio >1) with the daily probability of acquiring a pOXA-48 clone. Second, pre-existing colonisation with a pOXA-48 clone of a different species (lower part). This covariate measures how being previously colonised by a pOXA-48-carrying *E. coli* is associated with the daily probability of becoming colonized with a pOXA-48-carrying *K. pneumoniae* clone (Eco -> Kpn) and *vice versa* (Kpn -> Eco). We expect to see a positive association if within-patient transfer of pOXA-48 between different bacterial clones is important. Points represent posterior medians; thick grey lines represent the 80% CrI and thinner black lines represent the 95% CrI. (B) Number of previously uncolonised patients becoming colonised by a pOXA-48-carrying *K. pneumoniae* (top row) or *E. coli* (bottom row) as a function of the number of patients on the ward already colonised by a pOXA-48-carrying clone. (C) Number of R-GNOSIS study patients colonised by pOXA-48-carrying *K. pneumoniae* (Kpn) or *E. coli* (Eco) clones or both (co-colonised). For cocolonised patients, the colour code indicates whether *K. pneumoniae* or *E. coli* were isolated first or whether both species were simultaneously isolated from the same swab.

The baseline daily probabilities for becoming colonised with pOXA-48-carrying *K. pneumoniae* or *E. coli* were 0.1% (95% credible interval [CrI] 0.08%, 0.12%) and 0.04% (95% CrI, 0.02%, 0.05%), respectively (Supplementary Table 3). Results showed that the probability of acquisition of a pOXA-48-carrying *K. pneumoniae* was higher if other patients were already colonised with a pOXA-48-carrying clone in the wards of neurosurgery (Odds ratio [OR] 6.7, 95% CrI 2.5, 11.7) and pneumology (OR 2.7, 95% CrI 1.2, 4.6). In the wards of gastroenterology (OR 1.7, 95% CrI 0.4, 4.1) and urology (OR 0.6, 95% CrI 0.01,4.4) there were no clear effects. In contrast, the presence of other patients colonised by pOXA-48-carrying clones was not associated with the probability of acquiring a pOXA-48-carrying *E. coli* in the wards of neurosurgery (OR 0.23, 95% CrI 0.001,2.0) or urology (OR 0.4, 95% CrI 0.002, 2.7), and there was only weak evidence for a positive association in the wards of gastroenterology (OR 1.9, 95% CrI 0.4, 4.7) and pneumology (OR 2.6, 95% CrI 0.9, 4.5) (Figure 2A). This result suggested that *K. pneumoniae* is more important for between-patient transfer than *E. coli.*

The model also showed that prior colonisation with a pOXA-48-carrying clone of a different species was associated with a dramatic increase in the probability of acquiring a second pOXA48-carrying species (Figure 2A,C). This risk was high both when a patient was first colonised by *K. pneumoniae* (OR 23.3, 95% CrI 8.3, 53.4) and when initially colonized with *E. coli* (OR 15.8, 95% CrI 3.8, 42.7). This result underlines the potential importance of within-patient plasmid transfer in the disemination of pOXA-48, a role supported by the high frequency of co-colonised patients (Figure 2C). However, other explantions may be responsible for this observation, such as independent colonisation events of predisposed patients by different pOXA-48-carrying clones.

### Genomic analysis of pOXA-48-carrying enterobacteria

A key limitation of our epidemiological model is that it is based solely on species identification, which restricts the possibility of reconstructing the spread of specific clones and plasmids. To track within-patient and between-patient plasmid transfer at a higher level of resolution, we integrated genomic information by sequencing the genomes of the 171 pOXA-48-carrying isolates represented in Figure 1D. In line with previous studies^22,25^, the sequencing results revealed that a small subset of isolates initially identified as *K. pneumoniae* actually belonged to the species *Klebsiella quasipneumoniae* (n= 2) and *Klebsiella variicola* (n= 3) (Supplementary Figure 2).

We analysed the genetic relatedness of isolates belonging to *K. pneumoniae* and to *E. coli* separately by reconstructing the core genome phylogeny for each species (Figure 3). For *K. pneumoniae* (n= 103), most of the isolates belonged to a few high-risk sequence types: ST11 (n= 64), ST307 (n= 17), and ST15 (n= 9). In contrast, *E. coli* (n= 45) showed a more diverse population structure, with only one sequence type, ST10, comprising more than three isolates (n= 11).

**Figure 3.**
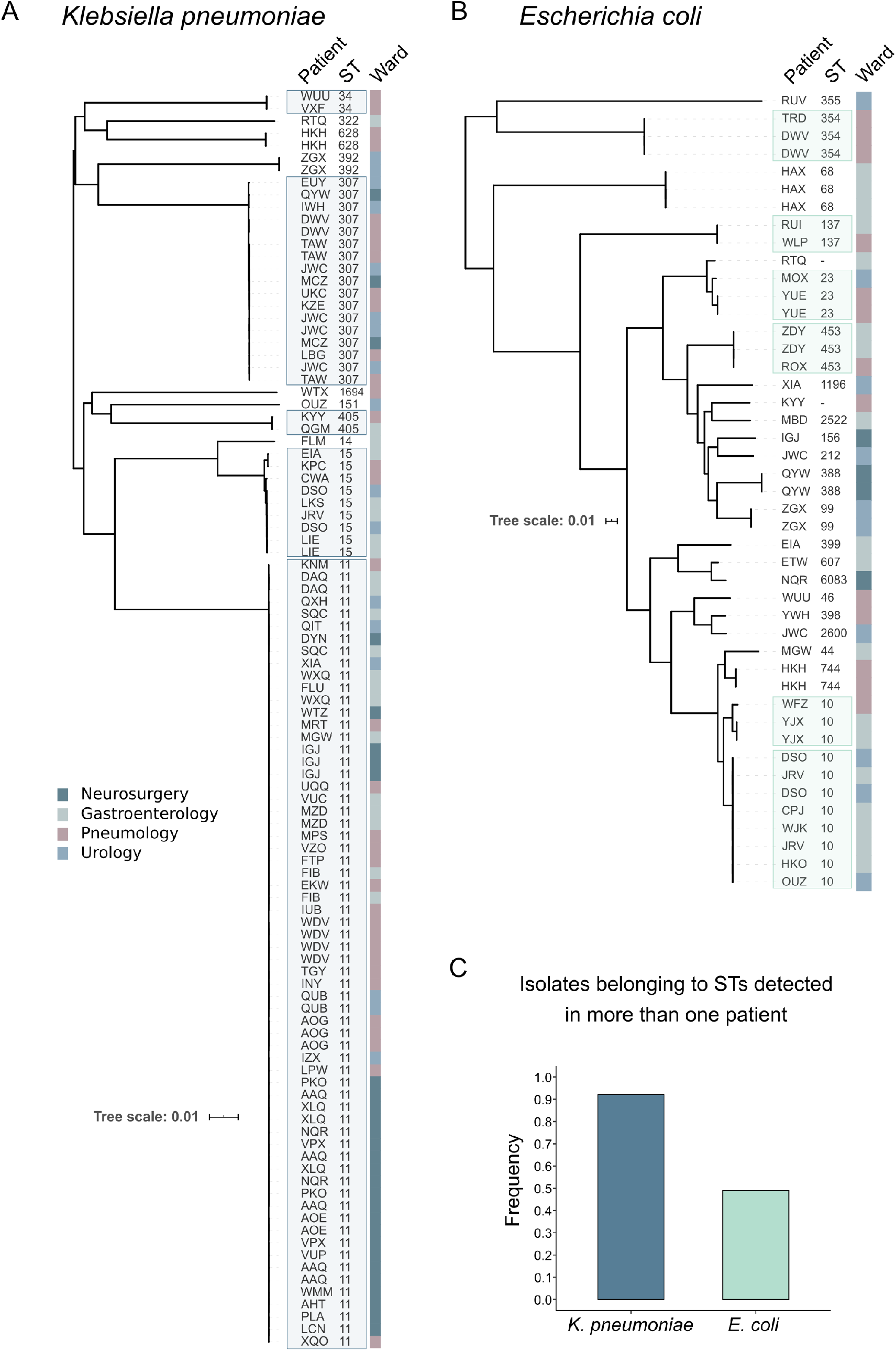
Phylogenetic analysis of pOXA-48-carrying *K. pneumoniae* and *E. coli.* Genetic relationships among (A) *K. pneumoniae* (n= 103) and (B) *E. coli* (n= 45) isolates carrying pOXA-48 and recovered during the R-GNOSIS study. Tree construction is based on polymorphisms in the core genome (scale: single nucleotide polymorphism [SNPs]/site). The columns to the right of the tree indicate patient code, isolate sequence type (ST), and the ward where the isolate was recovered (colour code in legend). Boxes with colour shading indicate recovery of isolates of the same sequence type (ST) from multiple patients in the hospital. (C) Frequency of pOXA-48-carrying *K. pneumoniae* and *E.* coli isolates belonging to STs detected in multiple patients.

We next considered the distribution of the different clonal groups (defined by the different STs) across colonised patients (Figure 3A,B). Most *K. pneumoniae* isolates belonged to STs present in more than one patient, whereas approximately half of *E. coli* isolates belonged to STs present in only one patient (Figure 3C). This result, together with the results of statistical analysis, suggested that a limited number of *K. pneumoniae* high-risk clones are responsible for most of the between-patient transfer events. However, we observed several cases of pOXA-48-carrying *E. coli* STs colonising different patients, suggesting that *E. coli* is also responsible for sporadic between-patient transmission events.

### Reconstruction of between-patient transfer dynamics of pOXA-48-carrying clones

To investigate the specific dissemination routes of pOXA-48-carrying clones, we integrated epidemiological and genomic data using SCOTTI^26^ (see methods). SCOTTI is a structured coalescent-based tool for reconstructing bacterial transmission, which accounts for bacterial diversity and evolution within hosts, non-sampled hosts, and multiple infections of the same host. We analysed the spread of the dominant *K. pneumoniae* and *E. coli* STs within and among the four wards under study (Figure 4, Supplementary Figures 3–10). As expected from the genomic data (Figure 3A), clones belonging to *K. pneumoniae* ST11 were responsible for most of the putative between-patient transmission events. The analysis attributed transmission events of pOXA-48-carrying ST11 on every single ward and even between wards, with neurosurgery being the ward with the highest frequency and probability of ST11 transmission (Figure 4), as suggested by the epidemiological model (Figure 2A). In light of these results, we investigated *K. pneumoniae* ST11/pOXA-48 epidemiology in the neurosurgery ward in more detail by looking at the spatiotemporal distribution of colonised patients (Supplementary Figure 11). The neurosurgery ward includes 11 rooms with 20 beds (9 double rooms and 2 single rooms). Of the 16 colonized patients, 6 had overlapping stays in the same room, suggesting that this room acted as a hotspot for *K. pneumoniae* ST11/pOXA-48 colonisation and transmission.

**Figure 4.**
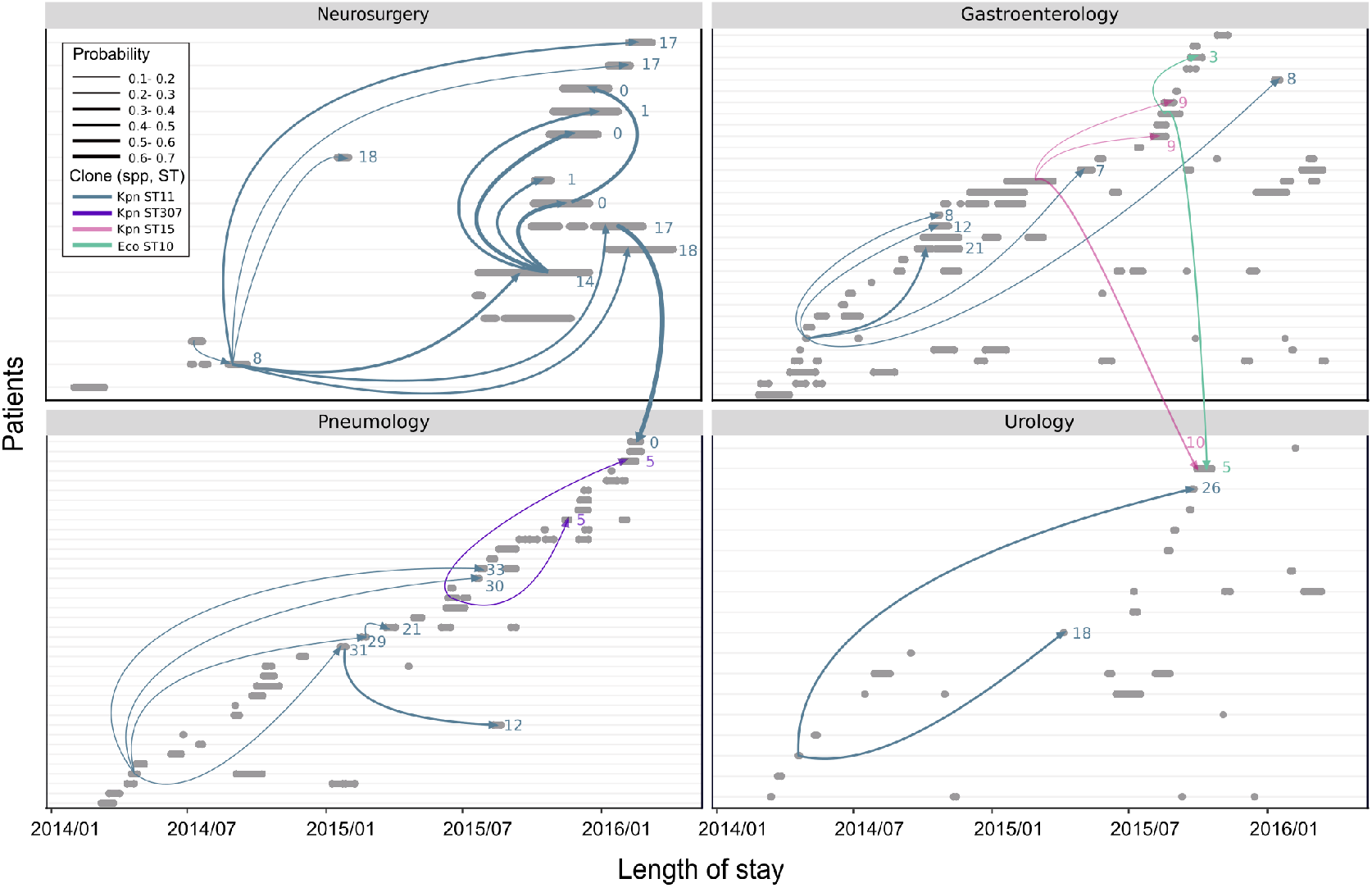
SCOTTI reconstruction of between-patient transfer of pOXA-48-carrying enterobacteria. The charts represent SCOTTI-attributed between-patient transfer events involving pOXA-48-carrying enterobacteria clones in the hospital, with individual panels representing the distribution of patients colonized by pOXA-48-carrying enterobacteria on the different wards. Each row represents an individual patient, and the grey segments represent the length of stay (from admission to discharge). Coloured arrows represent transmission events predicted by SCOTTI. Line colour indicates the clone responsible for the transmission event, and line thickness represents the probability of the SCOTTI-attributed transmission, as indicated in the legend: Kpn, *K. pneumonia;* Eco, *E. coli;* ST, sequence type. Numbers to the right of arrowheads indicate the number of SNPs differentiating the complete genomes of the clone pair involved in the putative transmission event.

SCOTTI also predicted transmission events mediated by three further pOXA-48-carrying clones. Two transmission events were attributed to *K. pneumoniae* ST307 in the pneumology ward and three more to *K. pneumoniae* ST15: two in gastroenterology and another one between the gastroenterology and urology wards. In line with the genomic results (Figure 3B), SCOTTI also attributed two between-patient transfer events to *E. coli* ST10, one on the gastroenterology ward and another one between the gastroenterology and urology wards (Figure 4).

### Genetic analysis of pOXA-48 confirms pervasive within-patient plasmid transfer

Our results suggest that the high frequency of patient colonisation by two plasmid-carrying species could be due to within-patient pOXA-48 transfer (Figures 1 and 2). However, although unlikely, another possibility is independent colonisation events involving different plasmid-carrying clones. To distinguish between these possibilities, we analysed the genetic sequence of plasmid pOXA-48 across all isolates with the aim of using specific genetic signatures in the plasmid to provide evidence for or against within-patient plasmid transfer. To increase the resolution of this analysis, we enriched the R-GNOSIS collection by recovering and sequencing the complete genomes of all the pOXA-48-carrying enterobacteria isolated from patients in our hospital since the plasmid was first reported in 2012 (Supplementary Table 1). In total, we determined and analysed the complete genomes of 250 strains, combining short-read and long-read sequencing technologies (see methods and Supplementary Figure 12).

The results showed that pOXA-48 is highly conserved (Figure 5A). The core plasmid sequence spanned more than 60 kb (>90% of plasmid sequence) in 218 of the 250 strains (Supplementary Table 1). Analysis of the core genome among these 218 plasmids revealed an identical sequence in 80% of them. In the remaining 20%, we detected a total 21 SNPs, with each plasmid presenting 1 or 2 SNPs compared with the most common variant (Figure 5A). This high degree of plasmid structural and sequence conservation and the strong link between pOXA-48 and the *bla*_oχA-48_ gene are important differences from previous analyses on the spread of plasmid-mediated carbapenemases such as KPC^8,9,27^.

**Figure 5.**
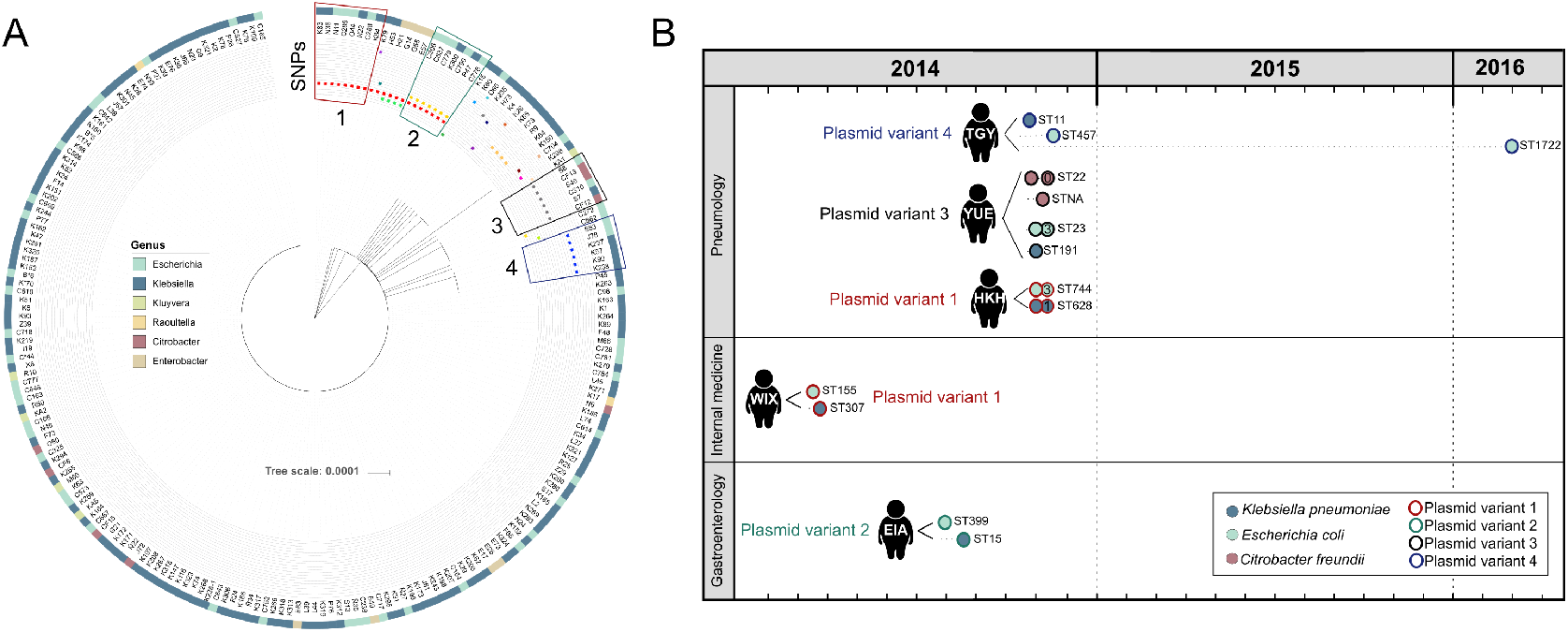
Pervasive within-patient pOXA-48 transfer. (A) Dendrogram constructed from the 21 polymorphisms present in the core region of plasmid pOXA-48. The outermost circle indicates plasmid-carrying clones genus according to the colour code in the legend, the second circle indicates the clone names, and the remaining circles indicate the presence of each plasmid SNP. Coloured boxes indicate the four plasmid variants carrying a ‘rare’ SNP present in clones of different species and used as genetic fingerprints. (B) Representation of patients colonized by clones carrying rare (traceable) plasmid variants. Patients are represented with their corresponding three-letter patient code. Circles represent clones isolated from the patient, with the fill colour indicating the bacterial species and the outline colour indicating the plasmid variant (see legend). The sequence type (ST) of each clone is indicated. Circles in the same row indicate different isolates of the same clone; the number inside the second circle indicates the number of SNPs accumulated in the complete genome relative to the first isolation.

Given the low plasmid-sequence variability, we could not track plasmid transmission using the same tools used for bacterial transmission. Instead, we monitored plasmid transfer by using the rare plasmid variants carrying specific core-region SNPs as genetic fingerprints (Figure 5). We focused on instances where the same traceable plasmid variant was present in different bacterial clones (belonging to different species). We considered that isolation of different bacterial species carrying the same rare plasmid variant from the same patient would be a very strong indicator of within-patient plasmid transfer. We found four examples in which the same rare plasmid variant was present in different bacterial species (Figure 5A). In all four, different species carrying the same plasmid variant were isolated from the same patient (Figure 5B). For example, plasmid variant 3 was detected in 6 bacterial isolates belonging to four clones (one *K. pneumoniae,* one *E. coli* and two *C. freundii),* and all of them were recovered from a single patient in the hospital (patient code YUE). Crucially, the chances of independent patient colonisation with the different bacterial clones carrying these rare plasmid variants are extremely low (variant 1, 6.4×10^-4^; variant 2, 8.9×10^-4^; variant 3, 1.1×10^-8^; variant 4, 2.1×10^-5^), confirming that these were within-patient plasmid transfer events.

### *High* in vitro *pOXA-48 conjugation rate*

Despite the limitations imposed by the sensitivity and frequency of the sampling method, the four selected pOXA-48 variants with core-region SNPs demonstrated pervasive within-patient plasmid transfer. However, the specific SNPs used as genetic fingerprints might affect the conjugation ability of the plasmid, which would make it impossible to generalize the results with these variants to the most common pOXA-48 variant. To exclude this possibility, we experimentally tested the conjugation rates of the most common pOXA-48 variant and the four traceable variants by introducing the plasmids independently into the *E. coli* strain J53 and determining the conjugation rate of each variant in this isogenic background (Figure 6, see methods). Although conjugation rates differed slightly (ANOVA; F= 2.9, df= 4, *P*= 0.037), they did not differ significantly between the traceable SNP plasmid variants and the most frequent variant (Tukey multiple comparisons of means, *P* > 0.3), indicating that all these plasmid variants have similar within-patient transfer ability. We therefore conclude that horizontal spread of pOXA-48 in the gut microbiota is the norm in colonized patients.

**Figure 6.**
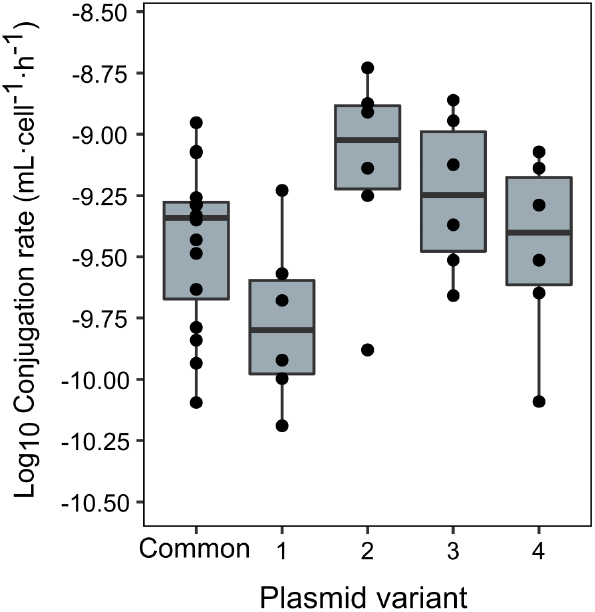
pOXA-48 conjugation rate. Conjugation rates of the most common pOXA-48 variant in the hospital (common, 12 biological replicates) and the four core-region SNP variants used to track within-patient plasmid transfer (6 biological replicates). Plasmid variant numbers correspond to those indicated in Figure 5. Horizontal lines inside boxes indicate median values, the upper and lower hinges correspond to the 25th and 75th percentiles, and whiskers extend to observations within 1.5 times the interquartile range.

Interestingly, and as previously reported^28^, the *in vitro* pOXA-48 conjugation rate was extremely high, with a median frequency of 0.3 transconjugants per donor after only one hour of mating (Supplementary Figure 13). The high pOXA-48 conjugation rate helps to explain the frequent within-patient plasmid transfer reported here.

## Discussion

CPE are emerging as one of the most concerning threats to public health worldwide^5^. Recent studies have highlighted the central relevance of hospitals as hotspots for the dissemination of CPE among patients and for the dissemination of the carbapenemase-encoding conjugative plasmids between enterobacteria clones^6–8^. In this study, we performed a high-resolution epidemiological analysis to uncover the dissemination routes of the carbapenemase-encoding plasmid pOXA-48 (both at the bacterial and plasmid levels). By integrating epidemiological and genomic data, we uncovered frequent between-patient bacterial transfer and pervasive within-patient plasmid transfer.

In light of our results, we propose that in-hospital pOXA-48 dissemination generally adheres to the following dynamics: high-risk pOXA-48-carrying enterobacteria clones, mainly *K. pneumoniae* ST11, spread among hospitalised patients, colonising their gut microbiota (Figures 2, 3 and 4). Once patients are colonised, the plasmid readily spreads through conjugation towards other resident members of the gut microbiota (enterobacteria such as *E. coli, C. freundii,* and *E. cloacae,* Figures 1 and 5). The plasmid’s high conjugation rate increases its chances of becoming established in the gut microbiota because, even if the invading nosocomial clone is eliminated, pOXA-48 can survive in a different bacterial host. Moreover, the frequent plasmid transfer provides a test bench for new bacterium-plasmid combinations, some of which may be particularly successful associations able to persist and even disseminate towards new human hosts^4^. An illustrative example of these general dynamics is the case of the patient carrying plasmid variant 4 (Figure 5B; patient code TGY). This patient was first colonised by *K. pneumoniae* ST11/pOXA-48 in October 2014, and 11 days later a pOXA-48-carrying *E. coli* strain was isolated from the same patient (ST457). During a new admission 17 months later, a different pOXA-48-carrying *E. coli* (ST1722) was recovered the patient’s gut microbiota. The pOXA-48 variant in all the clones carries a traceable SNP, confirming that the patient was colonized throughout the period by a pOXA-48-carrying enterobacteria, even though the plasmid had moved from its original *K. pneumoniae* ST11 host to *E. coli* clones in the gut microbiota.

Another interesting observation emerging from this study is that most of the events attributed to between-patient transmission originated from a small subset of patients (Figure 4). This result highlights the potential role of super-spreader patients in the nosocomial dissemination of CPE^29^. Unfortunately, given the small number of superspreader patients, we were not able to associate them to any particular epidemiological aspect, such as age or length of stay.

An important goal of this study is to inform new and improved intervention strategies aimed at controlling the spread of carbapenem resistance in hospitals. Our results can help in the design of interventions to control OXA-48 dissemination at two levels:

i. *Between-patient.* We have shown that the spread of pOXA-48-carrying enterobacteria between patients in the hospital is mainly mediated by high-risk clones commonly associated with nosocomial infections. These clones reside in hospital settings and are able to survive in the environment, creating stable reservoirs (often involving sinks^30–32^). Moreover, our results also showed that there are specific colonisation and transmission hotspots, such as individual rooms within wards (Supplementary Figure 11). We therefore propose that measures to detect and control environmental reservoirs and transfer hotspots could prevent between-patient OXA-48 dissemination. Such measures could represent a useful addition to the strategies based on patient surveillance and standard precautions already applied in hospitals, and could complement and improve the outcome of contact isolation measures^22^.
ii. *Within-patient.* A key finding of our study is the high prevalence of within-patient pOXA-48 transfer, which in turn helps to establish long-term pOXA-48 gut carriers. Preventing within-patient plasmid transfer and gut carriage is thus a particularly promising strategy for containing carbapenem resistance. This goal could be achieved either by blocking plasmid conjugation^33^ or, ideally, by specifically clearing pOXA-48 from the gut microbiota of patients by targeted decontamination. Decontamination strategies would aim to remove pOXA-48 plasmid or pOXA-48-carrying enterobacteria from carriers while preserving the integrity of the gut microbiota. New biotechnological advances are being made on this front. For example, CRISPR (clustered regularly interspaced short palindromic repeats) based technology can be used for targeted plasmid elimination^34^, and the new toxin–intein antimicrobials could be engineered to selectively remove pOXA-48-carrying clones from the microbiota^35^. Further work is urgently needed to tailor these emerging technologies into effective intervention strategies against the threat of plasmid-mediated carbapenemases.

## Methods

### Study design and data collection

We studied samples collected from patients admitted in a Spanish university hospital from March 4th, 2014, to March 31^st^, 2016, as part of an active surveillance-screening program for detecting ESBL/carbapenemase-carriers (R-GNOSIS-FP7-HEALTH-F3-2011-282512, www.r-gnosis.eu/)^18,22,23,36^. This study was approved by the Ramón y Cajal University Hospital Ethics Committee (Reference 251/13), which waived the need for informed consent from patients on the basis that the study was assessing ward-level effects and it was of minimal risk. This screening included a total of 28,089 samples from 9,275 patients admitted at 4 different wards (gastroenterology, neurosurgery, pneumology and urology) in the Ramon y Cajal University Hospital (Madrid, Spain). We used a randomly generated three-letters code for patient anonymization. Rectal samples were obtained from patients within 72 h of ward admission; weekly additional samples were recovered in patients hospitalised ≥7 days, and a final sample at discharge was obtained in those patients with a hospital stay ≥3 days (swabbing interval: gastroenterology, median 2 days, IQR 1,6 days; neurosurgery, median 3 days, IQR 1, 7 days; pneumology, median 2 days, IQR 1, 6 days; urology, median 1 day, IQR 1, 3 days). This protocol allowed us to obtain a time sequence for each patient in the hospital.

In this paper we have focused on the subset of patients colonised by pOXA-48-carrying enterobacteria within the R-GNOSIS project. Prevalence of colonisation by OXA-48-carrying enterobacteria among patients from 2014 to 2016 was 1.13% (105/9,275 patients). pOXA-48-carrying enterobacteria were the most frequent CPE in the hospital in this period, with 171 positive isolates (Supplementary Table 1). To better characterise pOXA-48 diversity and dissemination, we included in the within-patient pOXA-48 transfer analysis all the pOXA-48-carrying enterobacteria isolated from patients in our hospital since it was first reported in 2012. Specifically, we included 79 additional pOXA-48 carrying enterobacteria not included in the R-GNOSIS project (Supplementary Table 1).

### Bacterial characterisation

Samples were initially characterised as previously described, following the RGNOSIS protocol^23^. Briefly, swabs were plated on Chromo ID-ESBL and Chrom-CARB/OXA-48 selective agar media (BioMérieux, France) and bacterial colonies able to grow on these media were identified by MALDI-TOF MS (Bruker Daltonics, Germany). OXA-48 production was confirmed with KPC/MBL/OXA-48 Confirm Kit test (Rosco Diagnostica, Denmark). The MicroScan automated system (Beckman Coulter, CA, USA) was used for the antimicrobial susceptibility testing and the results were interpreted according to EUCAST guidelines (EUCAST breakpoint v7.1, www.eucast.org). Furthermore, specific *bla*_oχA-48_ resistance gene was identified by multiplex PCR^37^, and the PCR products were sequenced and compared with the GenBank database.

### Bacterial culture, DNA extraction, Illumina sequencing and PacBio sequencing

All pOXA-48-carrying enterobacteria isolates (n=250) were grown in Lysogeny broth (LB) medium at 37°C. Genomic DNA of all the strains was isolated using the Wizard genomic DNA purification kit (Promega, Madison, WI, USA), following manufacturer’s instructions. Whole genome sequencing was conducted at the Wellcome Trust Centre for Human Genetics (Oxford, UK), using the Illumina HiSeq4000 platform with 125 base pair (bp) paired-end reads. Furthermore, 2 isolates and 5 specific pOXA-48 plasmids (*K. pneumoniae* isolates k8 and k165, and plasmids from *K. pneumoniae* isolates k2, k164, k187, k236-1 and k273) were sequenced using the PacBio platform (The Norwegian Sequencing Centre; PacBio RSII platform using P5-C3 chemistry).

### Assembling and quality control (QC) analysis of sequence data

Trimmomatic v0.33^38^ was used to trim the Illumina sequence reads. SPAdes v3.9.0^39^ was used to generate *de novo* assemblies from the trimmed sequence reads with the -cov-cutoff flag set to ‘auto’. QUAST v4.6.0^40^ was used to generate assembly statistics. All the *de novo* assemblies reached enough quality including total size of 5–7Mb, the total number of contigs over 1 kb was lower than 200 and more than 90% of the assembly comprised contigs greater than 1 kb. Prokka v1.5^41^ was used to annotate the *de novo* assemblies with predicted genes.

### Phylogenetic analysis and identification of STs and clustering

Mash v2.0^42^ was used to determine distances between genomes using the raw sequence reads, and a phylogeny was constructed with mashtree v0.33^43^. For the analysis of the core genome (the set of homologous nucleotides present in all the isolates when mapped against the same reference) and the core sequence of the pOXA-48 plasmid, an alignment of the single nucleotide polymorphisms (SNPs) obtained with Snippy v2.5 (https://github.com/tseemann/snippy) was used to infer a phylogeny. A maximum-likelihood tree was generated using IQ-TREE with the feature of automated detection of the best evolutionary model^44^. All trees were visualised using the iTOL tool^45^. Recombination regions were identified with Gubbins^46^.

The seven-gene ST of all the isolates was determined using the multilocus sequence typing (MLST) tool (https://github.com/tseemann/mlst).

### Transmission mathematical modelling

Our statistical model was designed based on the premises established by Crellen T, *et al^24^.* The objective of our model is to estimate the daily probability of acquisition of a new pOXA-48-carrying enterobacteria by a patient in the hospital. Acquisition can occur through pOXA-48-carrying bacteria acquisition (between-patient transfer), or through pOXA-48 conjugation in the gut microbiota of the patient toward a new enterobacteria host (within-patient transfer).

We tracked all the pOXA-48-carrying enterobacteria identified in the hospital during the R-GNOSIS study period (Figure 1). This allows us to estimate and compare the acquisition of the most prevalent species, *K. pneumoniae* and *E. coli* independently. Each day in the ward, a patient can become colonised by a new pOXA-48-carrying *K. pneumoniae* or *E. coli*. However, as we lacked swabbing results from each day, the timing of new colonisation events with a pOXA-48-carrying clone are interval censored, and the analysis needs to account for this interval censoring^24^. If the probability of becoming colonised on day i for patient j is p_ij_, the probability of remaining uncolonized is (1-p_ij_). Therefore, in interval k for patient j consisting of N_kj_ days, the probability of remaining uncolonized is:

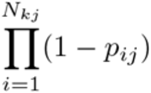

And the probability of becoming colonised (v_kj_) is the complement:

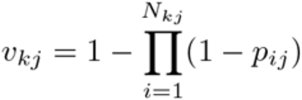

The outcome for patient j in interval k (X_kj_), is either that the patient acquired a new pOXA-48-carrying enterobacteria (X_kj_ =1) or did not (X_kj_ = 0). The likelihood is given by:

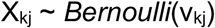

The daily probability of becoming colonised (p_ij_) is related by the logit link function to a linear function of covariates (π_ij_):

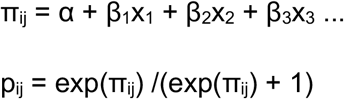

Where x represents a vector of predictors (data) and β is a vector of slopes (parameters) that are to be estimated. The β coefficient can be a single parameter, or permitted to vary by ward. The range of values and the prior distributions of the different parameters are described in Supplementary Table 2.

We developed and fitted models to study the probability of acquisition of pOXA-48-carrying *K. pneumoniae* and, separately, *E. coli.* We included the probability of *K. pneumoniae* and *E. coli* transferring the plasmid towards each other in the gut microbiota of colonised patients. To that end, we introduced as covariates the number of other patients already colonised by a pOXA-48-carrying enterobacteria each day, to study the risk of between-patient transfer (β coefficient), and if a patient was previously colonised with pOXA-48-carrying *E. coli* or *K. pneumoniae,* to study within patient pOXA-48-transmission (γ coefficient).

We considered five different transmission models to assess transmission of pOXA-48-carrying *K. pneumoniae* and *E. coli*:

1. Where the daily risk of acquiring pOXA-48-carrying *K. pneumoniae* and *E. coli* is constant (intercept only).
2. A constant term plus a between-patient transmission parameter β, where the explanatory variable (n_i_) is the number of patients colonised by pOXA-48 enterobacteria in the four wards.
3. As (2) but permitting the transmission parameter β to vary by ward (β_w_) and considering the number of patients colonised by a pOXA-48 enterobacteria in each ward (n_wi_).
4. As (2) but including a γ parameter for the within-patient transmission, and an explanatory variable (x_i_), which indicates if a patient had been previously colonised by a pOXA-48-carrying enterobacteria from a different species (yes, x_i_ = 1; no, x_i_ = 0).
5. As model (4) but permitting the transmission parameter β to vary by ward (β_w_) and considering the number of patients colonised by a pOXA-48 enterobacteria in each ward (n_wi_).

The probability of colonisation for individual j on day i for the respective models is calculated from:

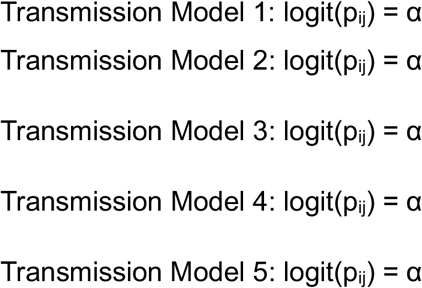

We fitted the statistical models using Hamiltonian Markov chain Monte Carlo in Stan (version 2.17.3) within the R environment (v. 3.4.3). Prior distributions were normal distributions using weakly informative priors^24^. Model comparison was performed with widely applicable information criterion (WAIC, Supplementary Table 2). The model that best fits our data is model number 5. We use 95% credible intervals (CrIs) to summarise uncertainty in posterior distributions. Daily probabilities calculated with model 5 are presented in Supplementary Table 3.

### Identification of transmission routes among patients

We applied SCOTTI^26^, a structured coalescent-based tool for reconstructing transmission, to the dominant *K. pneumoniae* and *E. coli* STs (with more than four isolates: *K. pneumoniae* ST11, ST15, ST307 and *E. coli* ST10), combining epidemiological and genomic data. We used the genome alignment, avoiding the recombination regions after the gubbins analysis^46^, as input to SCOTTI, together with the first date when an isolate was detected in a patient, and the start and end date of each patient’s infection risk period (Supplementary Table 1). Due to the possibility of transmission events between wards, we established a hierarchical analysis. First, we applied SCOTTI to the patients/genomes included in each ward to identify transmission routes within each ward, and second, we analysed the data of the 4 wards combined to identify additional transmission events between wards (Supplementary Figures 3–10).

### Identification of within-patient transmission routes of specific plasmid variants

In order to confirm within-patient plasmid transfer we studied specific pOXA-48 variants across the different isolates. The sequences belonging to pOXA-48 plasmid were mapped using the complete sequence of one of the plasmid sequenced by PacBio as reference (from *K. pneumoniae* k8), and the different variants and SNPs were identified using Snippy v2.5 (https://github.com/tseemann/snippy). We first analysed the degree of genetic variation in the plasmid among all the 250 bacterial clones. We compared the pOXA-48 variants sharing a core region of at least 60 kb (>90 % of the whole sequence, n= 218, Supplementary Table 1). We investigated cases where a variant of the plasmid carrying a “rare” traceable SNP is present in different clones (from different species). We found four plasmid variants present in different bacterial species and, in all cases, different species carrying the same plasmid variant were isolated from the same patient (Figure 5B). For those patients, we estimated the probability of those strains being acquired by independent subsequent transmissions events, assuming a random distribution of plasmid-carrying strains across patients. Analyses were performed using R (Version 3.4.2) (www.R-project.org).

### Conjugation assays

An initial conjugation round was performed to introduce pOXA-48 plasmids variants into *E. coli* J53^47^ (a sodium azide resistant laboratory mutant of *E. coli* K-12). pOXA-48-carrying wild type strains (donors) and *E. coli* J53 (recipient) were streaked from freezer stocks onto solid LB agar medium with selective pressure (ertapenem 0.5 μg/ml and sodium azide 100 μg/ml, respectively) and incubated overnight at 37°C. Three donor colonies and one recipient colony were independently inoculated in 2 ml of LB in 15-ml culture tubes and incubated for 1.5 h at 37°C and 225 rpm. After growth, donor and recipient cultures were collected by centrifugation (15 min, 1500 g) and cells were re-suspended in each tube with 300 μl of sterile NaCl 0.9%. Then, the suspensions were mixed in a 1:1 proportion, spotted onto solid LB medium and incubated at 37°C for 1.5 hours. Transconjugants were selected by streaking the conjugation mix on LB with ertapenem (0.5 μg/ml) and sodium azide (100 μg/ml). The transconjugants were verified by *bla*_oχA-48_ gene amplification by PCR as previously described^11^. For the isogenic conjugation experiments, the five different *E. coli* J53 carrying pOXA-48 plasmid variants acted as independent donors, and a chloramphenicol resistant version of J53 developed in our lab was used as the recipient strain. 6 colonies of each donor and recipient strains were independently inoculated in 2 ml of LB in 15-ml culture tubes and incubated overnight at 37 °C and 225 rpm. Each culture was used next day to inoculate 5 ml of LB in 50-ml culture tubes (1:100 dilution). After 1 hour of incubation at 37°C and 225 r.p.m, the pellets were collected by centrifugation (15 min, 1500 g) and cells were re-suspended in each tube with 300 μl of sterile NaCl 0.9%. Donor and recipient suspensions were mixed in a 1:1 proportion and plated on a sterile nitrocellulose filter (0.45 μm) on LB agar medium and incubated at 37°C for 1 hour. Simultaneously, each culture was plated on selecting agar for donors, recipient and transconjugants as controls (carbenicillin 100 μg/ml, chloramphenicol 50 μg/ml and a combination of both respectively). After 1 hour of incubation at 37°C, the filter contents were re-suspended in 2 ml of sterile NaCl 0.9%, diluted and plated on selective agar for donors, recipient and transconjugants. Transconjugants were verified by PCR as described above. Conjugation rates were determined using the end-point method^48,49^ (Figure 6), and the frequencies of transconjugants per donor were calculated from the same data (Supplementary Figure 13).

## Acknowledgements

This work was supported by the European Research Council under the European Union’s Horizon 2020 research and innovation programme (ERC grant agreement no. 757440-PLASREVOLUTION) and by the *Instituto de Salud Carlos III* (co-funded by European Development Regional Fund “a way to achieve Europe”) grants PI16-00860 and CP15-00012. The R-GNOSIS project received financial support from European Commission (grant R-GNOSIS-FP7-HEALTH-F3-2011-282512). RC acknowledges financial support from European Commission (R-GNOSIS) and *Plan Nacional de I+D+i2013–2016* and *Instituto de Salud Carlos III, Subdirección General de Redes y Centros de Investigación Cooperativa, Ministerio de Economía, Industria y Competitividad,* Spanish Network for Research in Infectious Diseases (REIPIRD16/0016/0011) co-financed by European Development Regional Fund “A way to achieve Europe” (ERDF), Operative program Intelligent Growth 2014–2020. BSC and TC acknowledge support from the UK Medical Research Council and Department for International Development [grant number MR/K006924/1], and BSC and PM acknowledge support under the framework of the JPIAMR - Joint Programming Initiative on Antimicrobial Resistance. ASM is supported by a Miguel Servet Fellowship (MS15-00012). JRB is a recipient of a Juan de la Cierva-Incorporación Fellowship (IJC2018-035146-I) co-funded by *Agencia Estatal de Investigación del Ministerio de Ciencia e Innovación*. MH-G was supported with a contract from *Instituto de Salud Carlos III,* Spain (iP-FIS program, ref. IFI14/00022). We thank the Oxford Genomics Centre at the Wellcome Centre for Human Genetics (funded by Wellcome Trust grant reference 203141/Z/16/Z) for the generation and initial processing of the sequencing data.

## Author contributions

ASM, RLS and BC conceived the study. RC designed and supervised sampling and collection of bacterial isolates. MHG, PRG collected the bacterial isolates and performed bacterial characterization. CDA and NLF collected the epidemiological data and performed preliminary analyses. R-GNOSIS WP5 Study Group designed sampling protocols and facilitated the training and capacity building for the collection of bacterial isolates and preliminary analyses. JdLF, JRB and CdlV performed the experimental work and analysed the results. RLS, BC, PM and TC performed the data analysis. ASM and RLS wrote the initial draft of the manuscript. ASM, RLS, JdlF, JRB, BC, PM, and TC contributed to the final version of the manuscript. All authors read and approved the manuscript.

## Competing interests

Authors declare no competing interests.

## Data availability

The sequences generated and analysed during the current study are available in the Sequence Read Archive (SRA) repository, BioProject ID: PRJNA626430, http://www.ncbi.nlm.nih.gov/bioproject/626430.

## Code availability

The code generated during the current study is available in GitHub, http://www.github.com/leonsampedro/transmission_stan_code.

## Supplementary tables

**Supplementary Table 2.** Transmission models to study the daily probability of acquisition of pOXA-48-carrying enterobacteria by hospitalised patients.

| Model | Parameters | Priors | *K. pneumoniae* |  | *E. coli* |  |
| --- | --- | --- | --- | --- | --- | --- |
|  |  |  | Posterior Median (95% CrI)<sup>1</sup> | WAIC<sup>2</sup> | Posterior Median (95% CrI)<sup>1</sup> | WAIC<sup>2</sup> |
| 1 | $\alpha$ (intercept) | Normal (0, 10) | -6.787 (-6.9797, -6.6019) | 1411 | -7.633 (-7.9330, -7.3640) | 705 |
| 2 | $\alpha$ (intercept)<br>$\beta$ (between-patient transfer) | Normal (0, 10)<br>Normal (0, 10) | -6.897 (-7.1103, -6.6990)<br>1.0611 (0.4981, 1.4857) | 1402 | -7.716 (-8.0609, -7.4121)<br>0.8333 (-0.7172, 1.5460) | 704 |
| 3 | $\alpha$ (intercept)<br>$\beta$ (between-patient transfer)<br>$M$ (beta mean)<br>$\Sigma$ (beta standard deviation) | Normal (0, 10)<br>Normal ($\mu$, $\sigma$)<br>Normal (0, 10)<br>Normal (0, 5) | -6.893 (-7.1098, -6.7027)<br>Varies by ward<br>0.6697 (-2.3853, 2.6142)<br>1.5789 (0.4050, 6.3881) | 1397 | -7.699 (-8.0255, -7.4054)<br>Varies by ward<br>0.0665 (-4.5453, 2.3773)<br>1.4629 (0.2166, 7.6397) | 706 |
| 4 | $\alpha$ (intercept)<br>$\beta$ (between-patient transfer)<br>$\gamma$ (within-patient transfer) | Normal (0, 10)<br>Normal (0, 10)<br>Normal (0, 10) | -6.919 (-7.1521, -6.7088)<br>1.0724 (0.3978, 1.4903)<br>2.7390 (1.2877, 3.7049) | 1394 | -7.827 (-8.1749, -7.5305)<br>0.6875 (-0.9421, 1.4782)<br>3.034 (2.0509, 3.8361) | 682 |
| 5 | $\alpha$ (intercept)<br>$\beta$ (between-patient transfer)<br>$M$ (beta mean)<br>$\Sigma$ (beta standard deviation)<br>$\gamma$ (within-patient transfer) | Normal (0, 10)<br>Normal ($\mu$, $\sigma$)<br>Normal (0, 10)<br>Normal (0, 5)<br>Normal (0, 10) | -6.924 (-7.1401, -6.7253)<br>Varies by ward<br>0.7198 (-3.0451, 2.9985)<br>1.5960 (0.3937, 6.7936)<br>2.7360 (1.2051, 3.7623) | 1388 | -7.836 (-8.1921, -7.5285)<br>Varies by ward<br>-0.2390 (-5.9522, 2.3138)<br>1.9315 (0.3361, 8.3896)<br>3.124 (2.1074, 3.9739) | 682 |

The table shows the parameters, prior and posterior distributions along with the WAIC (model comparison measure where lower values indicate a better fit to data). See methods for equations. Normal prior distributions show the mean and standard deviation respectively within brackets. <sup>1</sup> 95% Credible interval. <sup>2</sup> Widely applicable information criterion (WAIC).

**Supplementary Table 3.** Risk factors and the daily probability for acquisition of pOXA-48-carrying enterobacteria by hospitalised patients.

| Variable | Daily Probability of Acquisition<sup>1</sup><br>Posterior Median (95% CrI<sup>2</sup>) |  |
| --- | --- | --- |
|  | *K. pneumoniae* | *E. coli* |
| Baseline | 0.00098 (0.00079, 0.00120) | 0.00040 (0.00028, 0.00054) |
| Other patients colonised ($\beta$)<sup>3</sup> | | |
| Neurosurgery | 0.00650 (0.00276, 0.01089) | 0.00012 (2.39E-9, 0.00117) |
| Gastroenterology | 0.00163 (0.00034, 0.00385) | 0.00072 (4.86E-5, 0.00254) |
| Pneumology | 0.00268 (0.00115, 0.00445) | 0.00099 (0.00012, 0.00216) |
| Urology | 0.00056 (3.34E-7, 0.00381) | 0.00016 (5.88E-9, 0.00178) |
| Previously colonised patient ($\gamma$)<sup>4</sup> | 0.01489 (0.00330, 0.04009) | 0.00889 (0.00336, 0.01917) |

Daily probability of acquisition of pOXA-48-carrying *K. pneumoniae* or *E. coli* among patients admitted to the four different wards during the study period of the R-GNOSIS project. <sup>1</sup> Probability of acquisition is the posterior intercept plus the posterior coefficients transformed onto the probability scale. <sup>2</sup> Credible Interval of Posterior Distribution. <sup>3</sup> Risk of between-patient transfer: other patients already colonised by a pOXA-48-carrying enterobacteria each day, divided by ward. <sup>4</sup> Risk of within-patient transfer: patient previously colonised by a pOXA-48-carrying enterobacteria of the different species.

## Supplementary figures

**Supplementary Figure 1.**
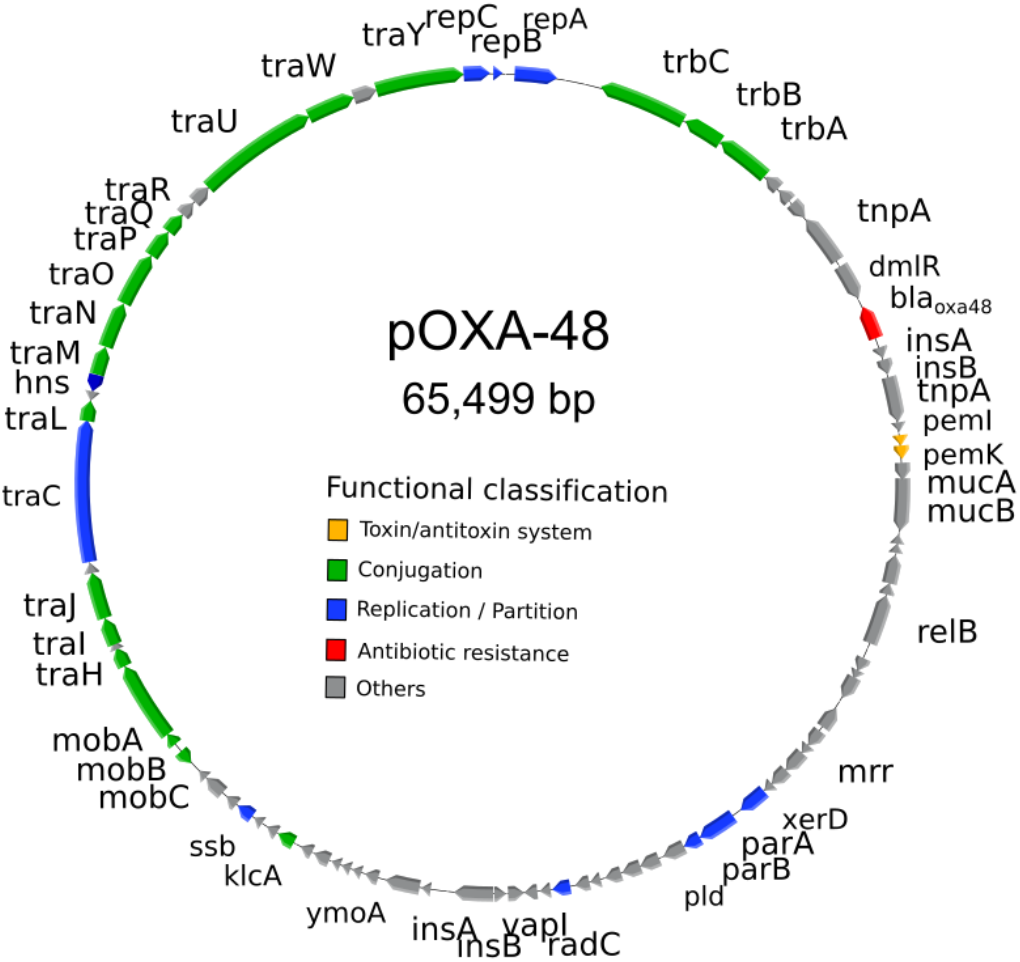
Plasmid pOXA-48. Schematic representation of plasmid pOXA-48. The reading frames for genes are shown as arrows, with the direction of transcription indicated by the arrowhead. Arrow colours indicate the functional classification of the gene (see legend). The *bla*_oχA-48_ gene is indicated in red.

**Supplementary Figure 2.**
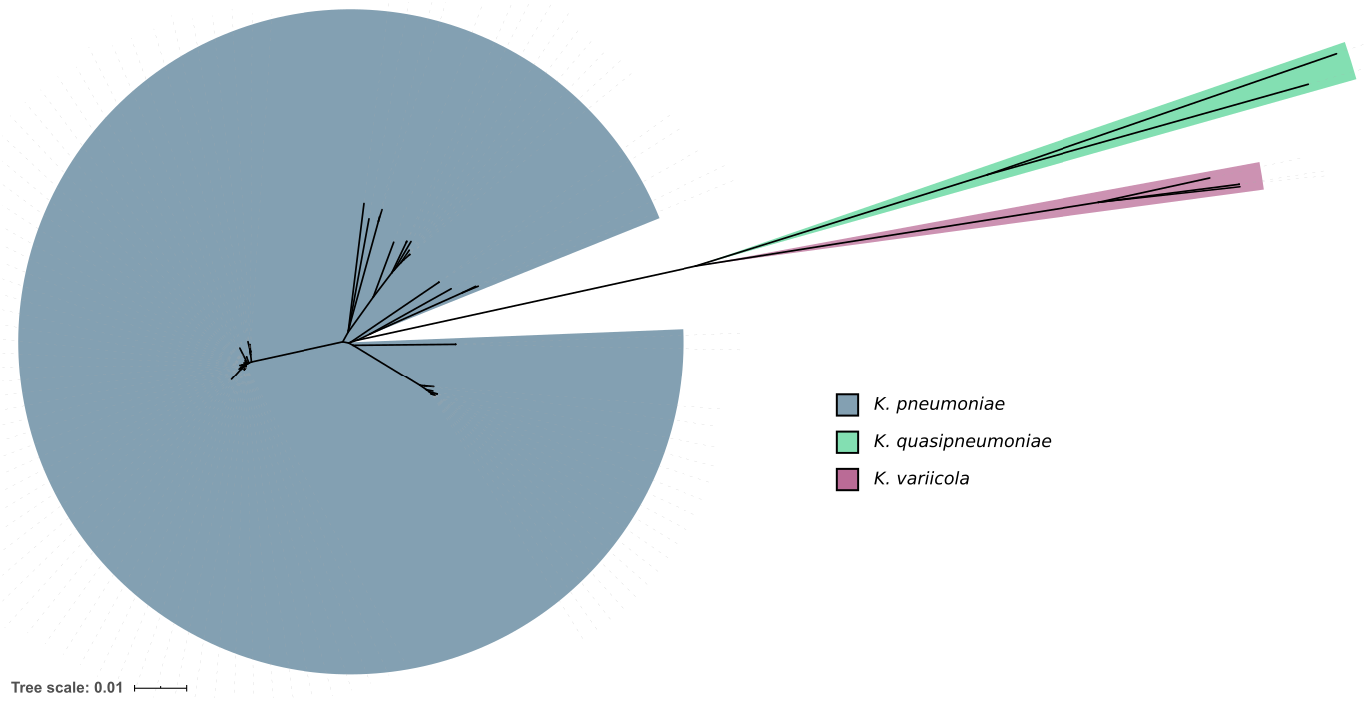
Phylogenetic analysis of isolates preliminary identified as *K. pneumoniae.* Unrooted phylogeny of 108 whole genome assemblies from the clones phenotypically identified as *K. pneumoniae.* Branch length gives the mash distance (a measure of k-mer similarity) between assemblies. Note the three distinct clusters, which are considered to be separate species (distance > 0.05): *K. pneumoniae* (n= 103), *K. quasipneumoniae* (n= 2) and *K. variicola* (n= 3).

**Supplementary Figure 3.**
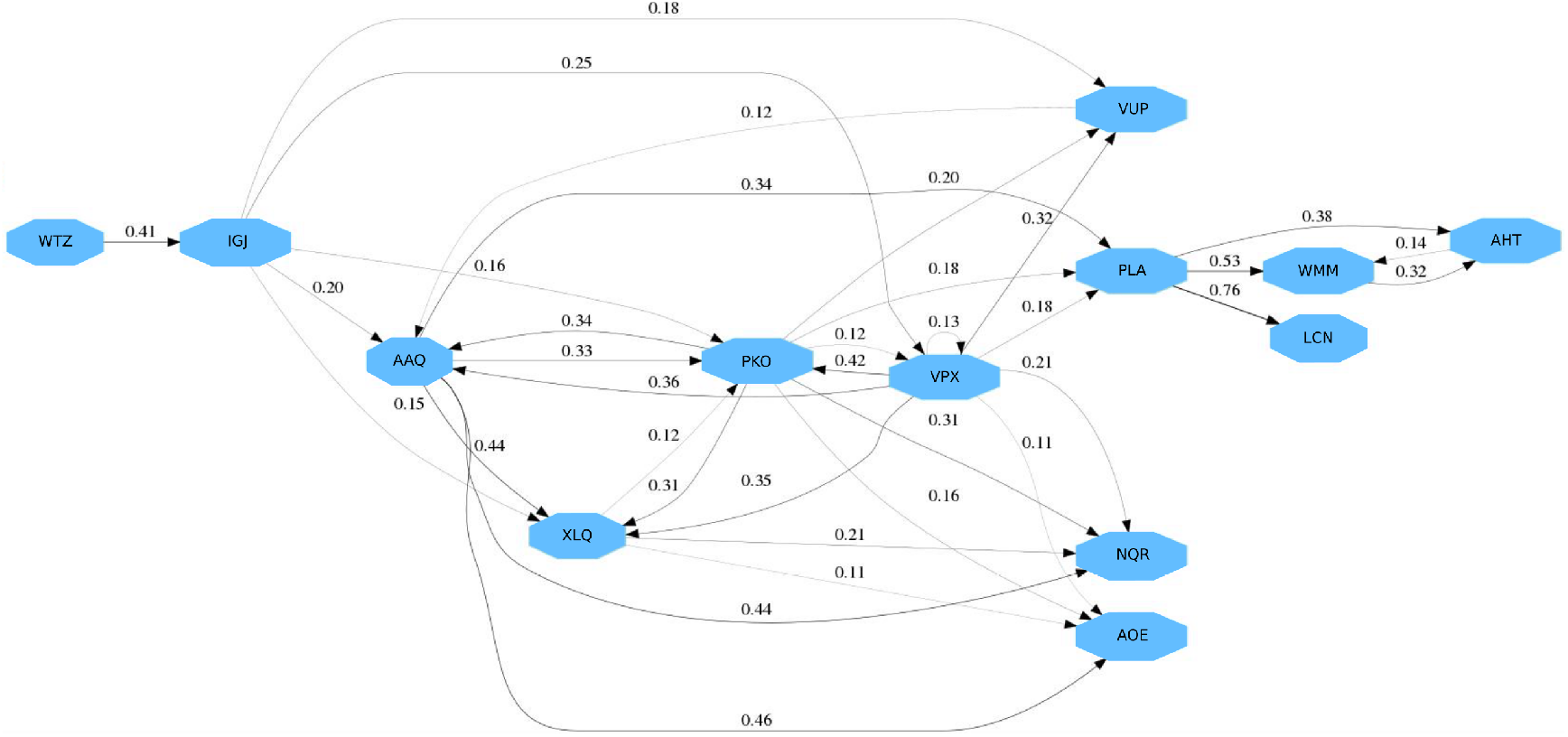
*K. pneumoniae* ST11 between-patient transfer dynamics in the neurosurgery ward. Transmission events of *K. pneumoniae* ST11 carrying plasmid pOXA-48 predicted by SCOTTI in the neurosurgery ward. Blue boxes represent patients, with patient codes indicated within the box. Lines represent the predicted between-patient transfer events, and the number above the lines indicate the probability of the transfer event.

**Supplementary Figure 4.**
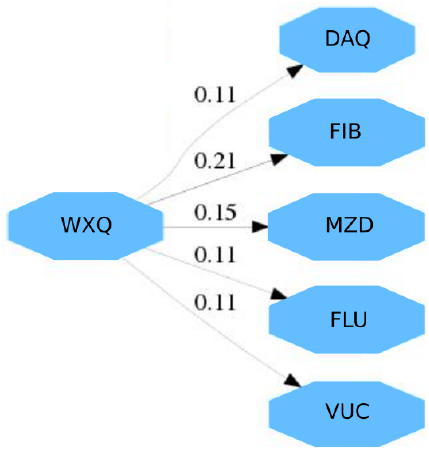
*K. pneumoniae* ST11 between-patient transfer dynamics in the gastroenterology ward. Transmission events of *K. pneumoniae* ST11 carrying plasmid pOXA-48 predicted by SCOTTI in the gastroenterology ward. Blue boxes represent patients, with patient codes indicated within the box. Lines represent the predicted between-patient transfer events, and the number above the lines indicate the probability of the transfer event.

**Supplementary Figure 5.**
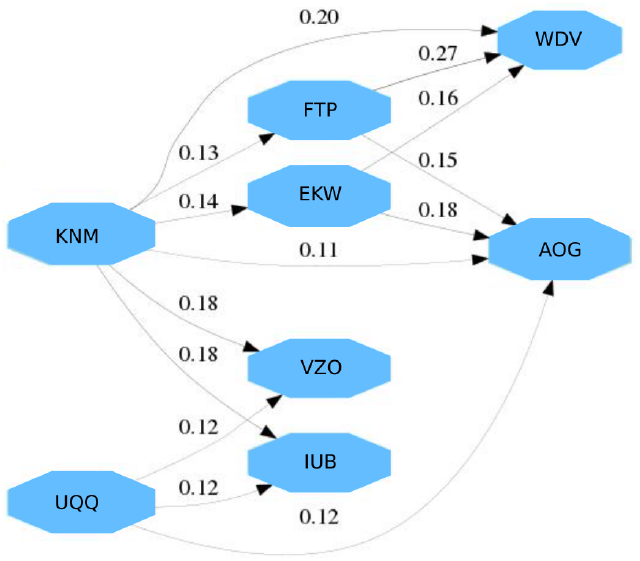
*K. pneumoniae* ST11 between-patient transfer dynamics in the pneumology ward. Transmission events of *K. pneumoniae* ST11 carrying plasmid pOXA-48 predicted by SCOTTI in the pneumology ward. Blue boxes represent patients, with patient codes indicated within the box. Lines represent the predicted between-patient transfer events, and the number above the lines indicate the probability of the transfer event.

**Supplementary Figure 6.**
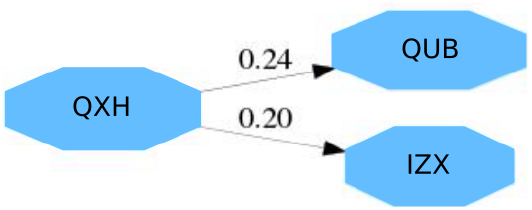
*K. pneumoniae* ST11 between-patient transfer dynamics in the urology ward. Transmission events of *K. pneumoniae* ST11 carrying plasmid pOXA-48 predicted by SCOTTI in the urology ward. Blue boxes represent patients, with patient codes indicated within the box. Lines represent the predicted between-patient transfer events, and the number above the lines indicate the probability of the transfer event.

**Supplementary Figure 7.**
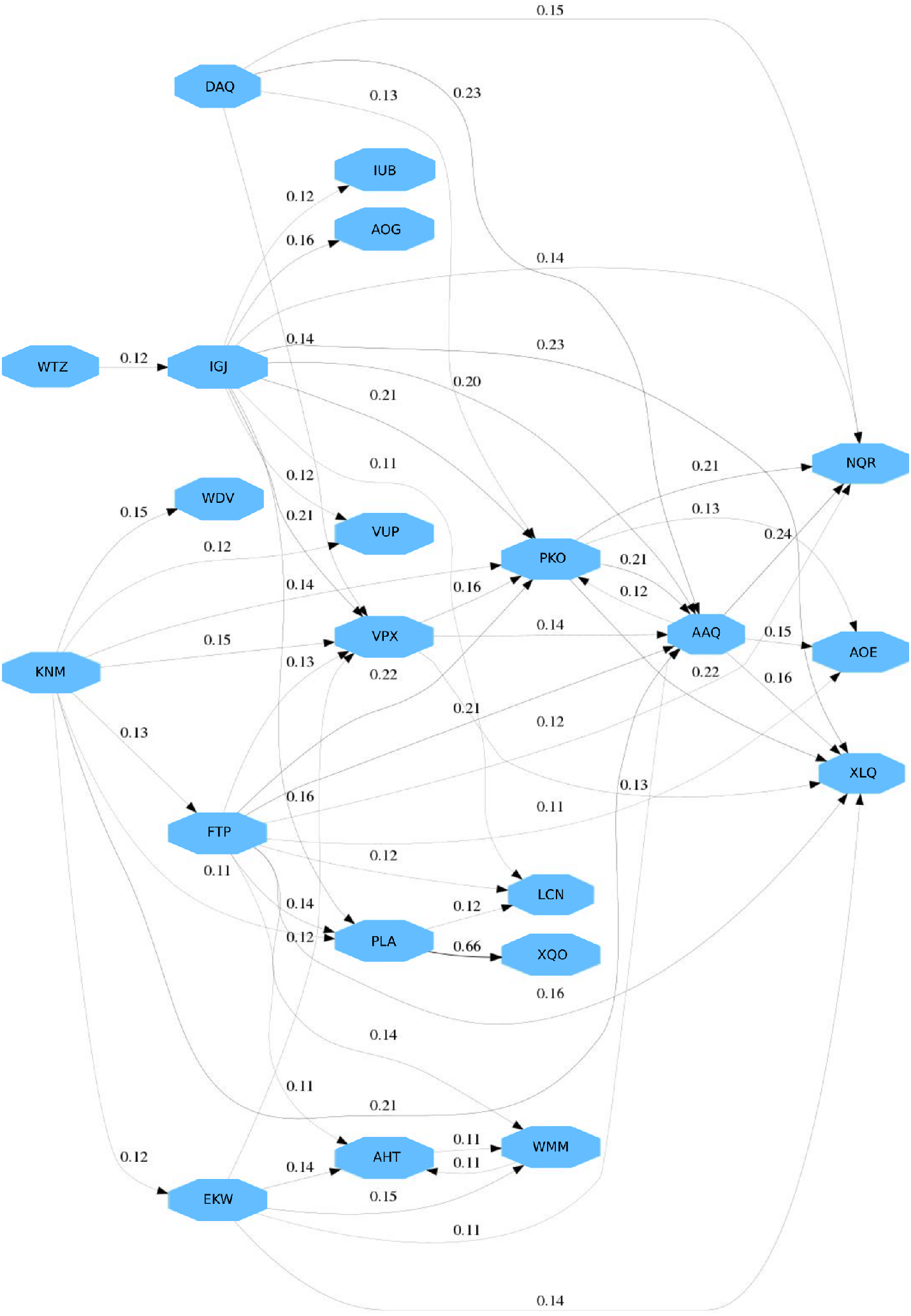
*K. pneumoniae* ST11 between-patient transfer dynamics across all the wards. Transmission events of *K. pneumoniae* ST11 carrying plasmid pOXA-48 predicted by SCOTTI when combining patients from the four wards. Blue boxes represent patients, with patient codes indicated within the box. Lines represent the predicted between-patient transfer events, and the number above the lines indicate the probability of the transfer event.

**Supplementary Figure 8.**
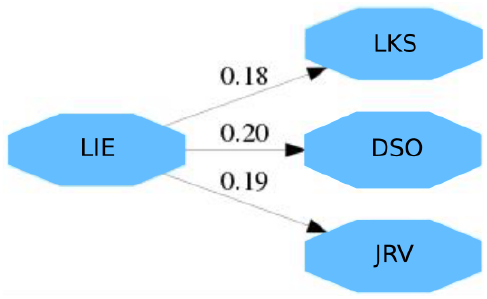
*K. pneumoniae* ST15 between-patient transfer dynamics across all the wards. Transmission events of *K. pneumoniae* ST15 carrying plasmid pOXA-48 predicted by SCOTTI when combining patients from the four wards. Blue boxes represent patients, with patient codes indicated within the box. Lines represent the predicted between-patient transfer events, and the number above the lines indicate the probability of the transfer event.

**Supplementary Figure 9.**
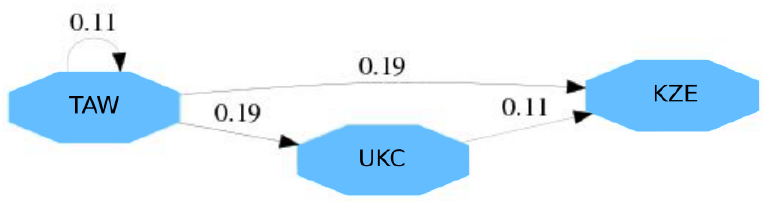
*K. pneumoniae* ST307 between-patient transfer dynamics in the pneumology ward. Transmission events of *K. pneumoniae* ST307 carrying plasmid pOXA-48 predicted by SCOTTI in the pneumology ward. Blue boxes represent patients, with patient codes indicated within the box. Lines represent the predicted between-patient transfer events, and the number above the lines indicate the probability of the transfer event.

**Supplementary Figure 10.**
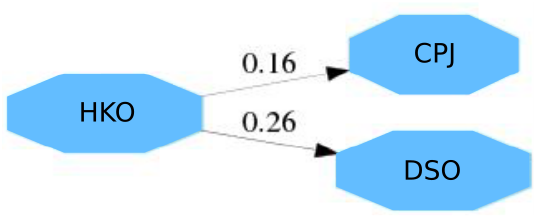
*E. coli* ST10 between-patient transfer dynamics across all the wards. Transmission events of *E. coli* ST10 carrying plasmid pOXA-48 predicted by SCOTTI when combining patients from the four wards. Blue boxes represent patients, with patient codes indicated within the box. Lines represent the predicted between-patient transfer events, and the number above the lines indicate the probability of the transfer event.

**Supplementary Figure 11.**
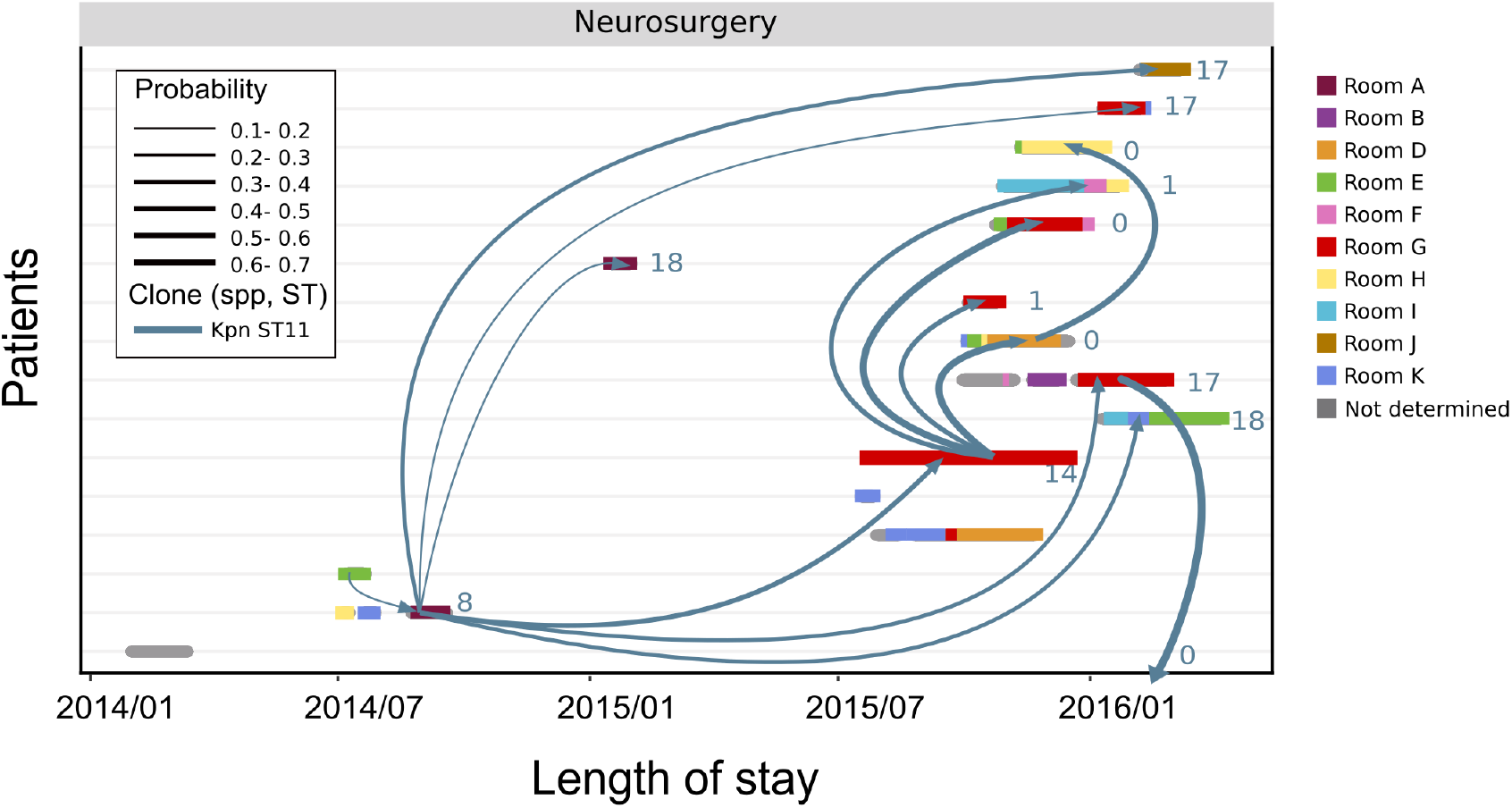
Spatiotemporal distribution of patients colonised by *K. pneumoniae* ST11 in the neurosurgery ward. Distribution of patients colonised by pOXA-48-carrying *K. pneumoniae* ST11 in the neurosurgery ward. Each row represents a patient and the colour segments represent the length of stay in the hospital (from admission to discharge). The colours of the segments represent the different rooms within the ward (see legend). Arrows represent transmission events predicted by SCOTTI. Line thickness represents the probability of the transmission predicted by SCOTTI. The number to the right of the arrowhead indicates the number of SNPs between the complete genomes of the pair of clones involved in the putative transmission event. Note that 6 out of 16 patients shared room G in overlapping stays.

**Supplementary Figure 12.**
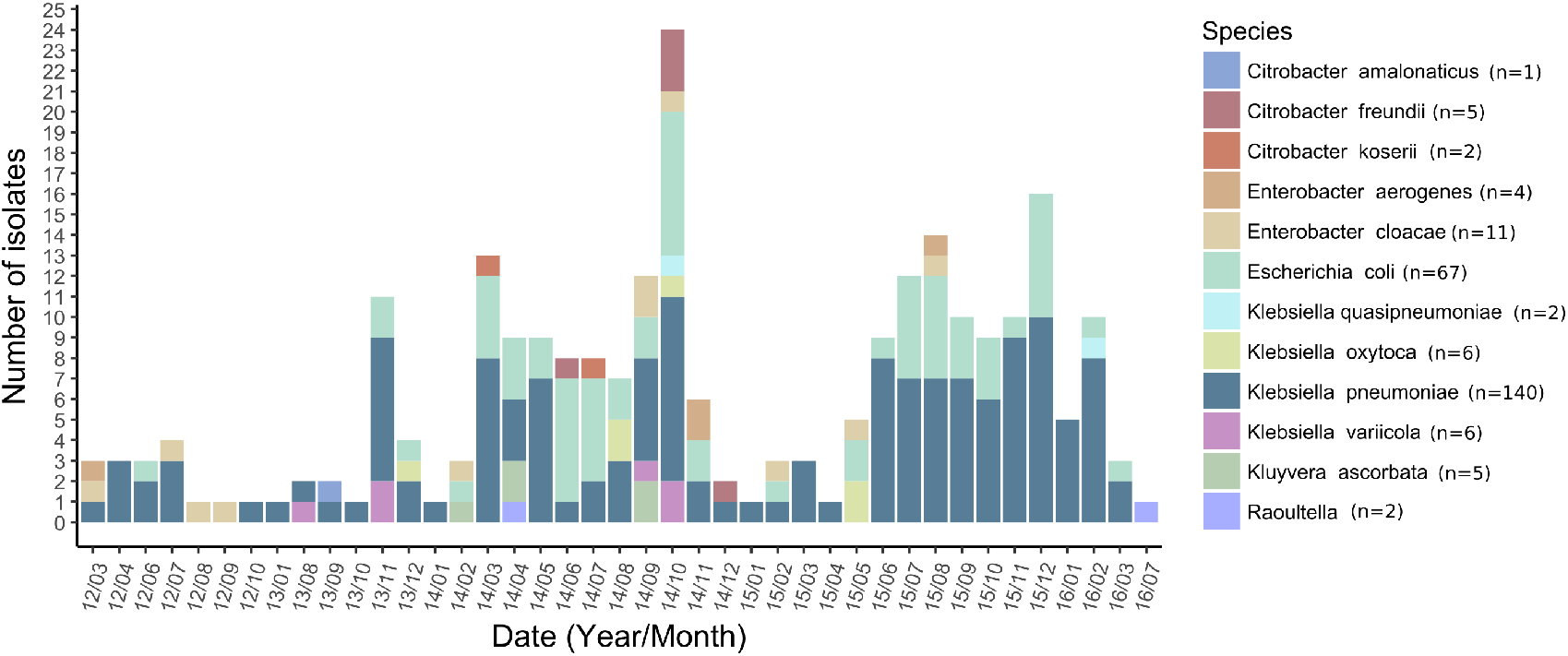
pOXA-48-carrying enterobacteria analysed in this study. Representation of the 250 pOXA-48-carrying clones isolated in the hospital from the first description till the end of the study period. The colour code indicates the species of the pOXA-48-carrying enterobacteria as indicated in the legend.

**Supplementary Figure 13.**
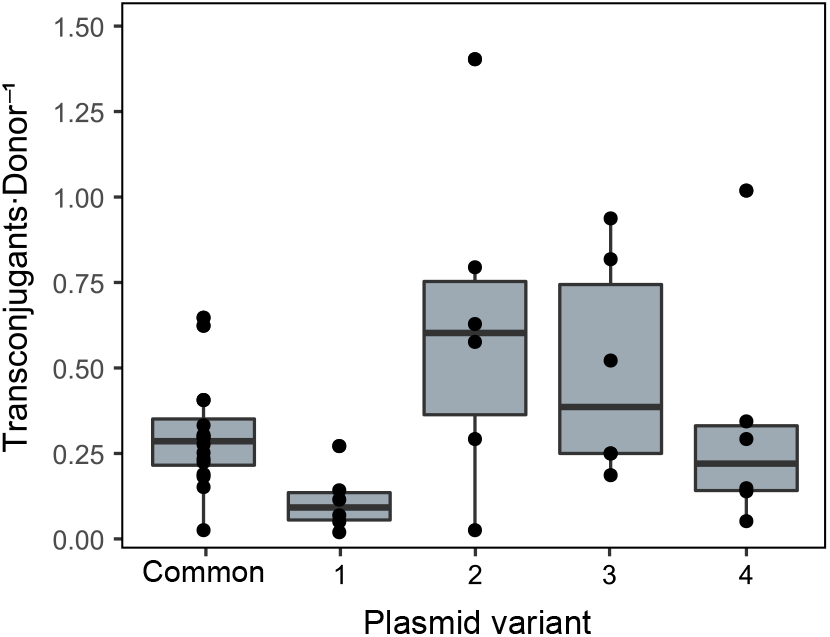
Frequency of conjugation of plasmid pOXA-48. Conjugation frequencies (transconjugants per donor) of the most common pOXA-48 variant in the hospital (common, n= 12 biological replicates) and the four variants with SNPs in the core region used to track within-patient plasmid transfer (n= 6 biological replicates). Plasmid variant numbers correspond to those indicated in Figure 5. The line inside the box marks the median. The upper and lower hinges correspond to the 25th and 75th percentiles and whiskers extend to observations within 1.5 times the interquartile range. The data presented here is the same as in figure 6, but represented as conjugation frequency instead of rate.

## Notes

http://www.ncbi.nlm.nih.gov/bioproject/626430

http://www.github.com/leonsampedro/transmission_stan_code

### Competing Interest Statement

The authors have declared no competing interest.

